# Comparison of transthoracic ultrasonography, computer-assisted lung auscultation, and transtracheal wash cytology in stocker cattle with bovine respiratory disease

**DOI:** 10.1101/2023.08.16.553541

**Authors:** Matthew A. Scott, Amelia R. Woolums, Hannah F. Carter, Robert W. Wills, Camilo Bulla, Brandi B. Karisch

**Author notes:** Correspondence: MAS,; ARW. These authors contributed equally to this work (Matthew A. Scott, Amelia R. Woolums).

## Abstract

Field methods to diagnose bovine respiratory disease (BRD) do not accurately identify airway inflammation and lack clinical sensitivity. New diagnostic modalities, such as thoracic ultrasound (TU), computer-assisted lung auscultation (CALA), and transtracheal wash (TTW), have recently emerged which may deliver accurate diagnosis and prediction of BRD in clinical settings. Therefore, we sought to compare TU, CALA, and TTW fluid cytologic assessment in stocker cattle at risk for BRD in a pilot study. We enrolled 17 high-risk mixed-breed beef steers, sampled 10 and 21 days after arrival and conventional management, in a pilot cross-sectional observational study. Cattle were examined daily for 82 days for clinical BRD. On day 10, 16 cattle received CALA, and 10 and 8 of these received TU and TTW, respectively. On day 21, 12 cattle received CALA and TTW, and 10 received TU. CALA was scored as 1-5. Lung consolidation and/or comet tails were evaluated by TU. TTW was evaluated by 200-cell differential count, with inflammation defined as >20% neutrophils. Relationships between each diagnostic test, and between diagnostic tests and clinical BRD, were evaluated by logistic regression (P<0.10). Fourteen cattle were treated for BRD. CALA scores ranged 1-3; three cattle had lung consolidation. On day 10, 5 of 6 cattle previously treated for BRD and 0 of 3 not treated had >20% TTW neutrophils. On day 21, 5 of 9 treated cattle and 1 of 3 not treated had >20% TTW neutrophils. No significant relationship between CALA, TU, and TTW inflammation existed. TTW inflammation was associated with BRD diagnosis (P=0.0586). CALA and TU results were unrelated to TTW inflammation. Cytologic assessment of TTW may improve antemortem diagnosis of BRD.

## 1. Introduction

Despite decades of research, bovine respiratory disease (BRD) remains the leading cause of morbidity and mortality in post-weaned beef cattle. In the United States, BRD is estimated to cause approximately $1 billion in production loss per year [1,2]. The efficacy of BRD control is limited by an inadequate diagnosis of the disease. While new diagnostic technologies are emerging, no diagnostic gold standard exists, and clinical assessment relies heavily on nonspecific clinical signs, such as depressed attitude and anorexia. Reliance on clinical signs for BRD diagnosis is suboptimal, as the sensitivity and specificity of clinical diagnosis are estimated to be 29-62% and 63-92%, respectively [3,4]. An improved means of antemortem BRD diagnosis is needed to advance therapeutic and preventive management strategies.

New diagnostic strategies are being tested to advance BRD diagnostic and prognostic accuracies with economic practicality. Two strategies of particular interest are transthoracic ultrasonography (TU) [5-8] and computer-assisted lung auscultation (CALA) [9-11]. Lung consolidation identified by TU has been associated with an increased risk of mortality [12] or relapse after treatment [8] in feedlot steers. Results of CALA have also been associated with increased risk of mortality in feedlot cattle [13], and the method has recently been used to select cattle for metaphylactic antimicrobial therapy [11]. However, either of these methods may fail to identify cattle in the early stages of BRD before lung consolidation or abnormal lung sounds have developed.

Cytological evaluation of the transtracheal wash (TTW), which may be a more sensitive means of identifying BRD onset and development than TU or CALA, remains understudied in bovine medicine. The technique yields a sample of tracheal secretions that represents a composite sample of lower airway secretions [14-16]. Cytological assessment of a TTW sample to identify the proportion of neutrophils and mononuclear cells, the relative amount of mucus, and the presence of bacteria, is commonly used to support a diagnosis of infectious or allergic lower airway disease in horses [15,16]. While the technique is likely to be too time-consuming and costly for routine use in commercial feedlot or stocker settings, in research applications it may help define more sensitive methods of diagnosis that are more accurate than clinical diagnosis and also field-expedient.

The objective of this pilot study was to compare the results of TU, CALA, and TTW cytology in post-weaned beef cattle at high risk for developing BRD. A second objective was to assess the association between the results of these three diagnostic tests and a recent clinical diagnosis of BRD. Our overarching hypotheses were that the results of the three tests would demonstrate moderate to high agreement and that abnormal results identified by these three diagnostic modalities would be positively related to a recent diagnosis of BRD.

## 2. Materials and Methods

### 2.1. Animal Enrollment and Processing

This report adheres to the ARRIVE 2.0 guidelines for reporting animal research [17]. All animal use and procedures were approved by the Mississippi State University Animal Care and Use Committee (IACUC protocol #17-120) and carried out in accordance with relevant IACUC and agency guidelines and regulations. Commercial crossbreed bulls (n=17; mean = 220 kg; range = 188-246 kg) were purchased from regional cattle auction markets within regional cattle auction markets within Mississippi and housed at the H. H. Leveck Animal Research Center at Mississippi State University. This experiment was performed on a subset of extra cattle from a larger study examining the effect of on-arrival vaccination and deworming on health and performance outcomes [18]. Upon arrival, cattle were weighed, assessed for elevated rectal temperature, and surgically castrated by a project veterinarian. All animals were given identification ear tags and tested for persistent infection of bovine viral diarrhea virus (BVDV) via ear notch antigen capture ELISA; no BVDV-positive calves were identified. Cattle did not receive mass medication at arrival, but they were vaccinated with a modified attenuated multivalent respiratory viral vaccine (Pyramid 5, Boehringer Ingelheim Vetmedica; Duluth, GA, USA) containing bovine herpesvirus-1 (BHV-1), bovine respiratory syncytial virus (BRSV), parainfluenza type 3 virus (PI3V), and bovine viral diarrhea virus types 1 and 2 (BVDV 1 + 2), and a multivalent clostridial bacterin-toxoid containing *Clostridium chauvoei, septicum, novyi, sordelli*, and *perfringens* Types C and D (Vision 7 with SPUR, Merck Animal Health; Madison, NJ, USA). The cattle were also treated with fenbendazole (SafeGuard, Merck Animal Health; Madison, NJ, USA) at 10 mg/kg PO and levamisole (Prohibit, Agri Laboratories; Saint Joseph, MO, USA) at 8 mg/kg PO for presumed gastrointestinal parasitism. The cattle were managed in one 2.5-acre grass paddock with mineral-mix provision over an 82-day period.

### 2.2. Study Design

This study was a pilot cross-sectional observational study, with observations made on day 10 and day 21 after arrival. Over the 82-day post-arrival period, the cattle were assessed daily by trained staff for clinical signs of BRD or other health problems. Clinical signs of BRD were assigned severity scores of 0-4 based on an approach adapted from that described by Step and colleagues [19]. Animals diagnosed with clinical BRD (score of 1 or 2 with a rectal temperature greater than 40° C or score of 3 or 4 regardless of temperature) were treated with antimicrobials as previously described [20]. Briefly, the first BRD treatment was with ceftiofur crystalline free acid (Excede, Zoetis; Parsippany, NJ, USA) at 6.6 mg ceftiofur equivalents/kg SC at the base of the ear. The cattle meeting criteria for BRD treatment eight days or greater after the first treatment were treated with florfenicol (Nuflor, Merck Animal Health; Madison, NJ, USA) at 40 mg/kg SC. The cattle meeting criteria for BRD treatment 5 days or greater after the second treatment were treated with oxytetracycline (300 mg/ml) (Noromycin 300 LA, Norbrook; Overland Park, KS, USA) at 20 mg/kg SC. Cattle showing signs of BRD after the third treatment were not treated again with antimicrobials but were evaluated by the project veterinarian to determine if they met the criteria for euthanasia. The cattle were weighed at arrival and on days 12, 26, and 82 post-arrival. Body weights, rectal temperature, treatments, and clinical BRD scores were recorded for each animal in an Excel spreadsheet (Microsoft Corporation; Redmond, WA, USA). On days 10 and 21 post-arrival, cattle were individually restrained within a portable manual dual-sided adjustable chute and were evaluated with CALA, TU, and TTW.

### 2.3. Computer-Assisted Lung Auscultation (CALA)

Computer-assisted airway auscultation (CALA) was performed with the Whisper® electronic stethoscope and BRD detection system software per manufacturer instruction (Geissler Corporation; Minneapolis, MN, USA). Briefly, the stethoscope was placed over the 5th intercostal space of the right thoracic wall, where respiratory sounds were recorded over a period of 8 seconds. Recorded audio was transferred to a laptop workstation within 3 meters of the stethoscope, where the software program spectrographically displayed the recorded audio and assigned ordinal scores of 1-5 (normal to severe) per Whisper® scoring software implementation. Although the manufacturer’s instructions indicated that the CALA should be assessed only on the right side of the thorax, the left side was also assessed in order to determine whether assessing both sides improved the test diagnostic performance.

### 2.4. Transthoracic Ultrasonographic (TU) Evaluation

The 3rd to 9th intercostal spaces on both left and right thoracic walls were scanned with an 8.5 MHz linear probe (IBEX PRO; EI Medical; Loveland, Co, USA); the imaging procedure was similar to the procedure described by Buczinski and colleagues [21]. Briefly, comet-tail artifacts and consolidation were recorded over the entire visible surface of the lung on both the right and left sides. Comet tail artifacts [22] were scored 0-3 (absent-to-severe). Lung consolidation was scored into two separate categories: shallow (> 2 cm) and deep (> 5 cm); cattle were scored into each of the two categories with either a 0 (absent) or 1 (present). Assessments were made by a single experienced examiner who was unaware of the treatment status of the cattle at the time of the examination.

### 2.5. Transtracheal Wash (TTW) Cytology Evaluation

Cattle sampled for TTW were restrained with the head elevated. A 15 cm2 area of ventral trachea at the junction between the proximal one-third and middle one-third of the palpable trachea was clipped for sterile site preparation, utilizing aseptic scrubbing with 2% chlorhexidine and 70% isopropanol. The area was then locally anesthetized with 2 mL of 2% lidocaine. The trachea was manually fixed, and a 15-g cannula and 48-cm catheter (MILA International; Florence, KY, USA) were placed for TTW sample collection. Thirty mL of sterile 0.9% isotonic saline was instilled and immediately withdrawn. The subsequent wash sample was transferred to an EDTA tube and stored on ice for the preparation of slides within 12 hours of collection for later cytological evaluation. Samples were prepared for both Cytospin centrifugation and direct cytology smears and then Wright-stained for cytologic evaluation. Cytology samples were microscopically assessed via a 200-count cell differential by a single experienced laboratory examiner who was unaware of the treatment status of the cattle at the time of the examination; direct smear preparations were counted if an adequate number of cells were present to complete a 200-cell differential count, and the Cytospin preparations were used if the direct smear was not adequately cellular. Cells counted included airway epithelial cells, macrophages, lymphocytes, neutrophils, eosinophils, basophils, and multinucleated cells. Slides were also scored semi-quantitatively for mucus on a scale of 0-2 and the presence of blood on a scale of 0-2. The TTW was defined as an inflammatory response if the percentage of neutrophils counted was 20% or higher. As guidelines for interpretation of bovine TTW have not, to our knowledge, been reported, this definition was selected based on guidelines for interpretation of equine TTW [23].

### 2.6. Data Analysis

The relationship between CALA score, TU score, and the presence of TTW inflammation (20% neutrophils or greater) on day 10 and day 21, and the relationship between clinical BRD diagnosis and CALA score, TU score, and TTW inflammation, was evaluated by logistic regression with SAS v9.4 (SAS Institute; Cary, NC, USA). For the TU comet tail score, the scores for the right and left sides of the thorax were added, and the sum of the scores was used in the analysis. The association between TTW inflammation and treatment for BRD within 0, 1, 2, 3, 4, 5, 6, or 7 days of sampling was assessed by logistic regression using PROC LOGISTIC with the EXACT function due to data scarcity; separate models were utilized for 1 to 7 days before and after BRD treatment. All statistical analyses were determined to be significant with an a priori P < 0.10. Descriptive statistics for TTW cytology were performed with Excel (v2306 Build 16+, Microsoft Corporation; Redmond, WA, USA).

## 3. Results

### 3.1. Cattle health and weight gain

The descriptive data for the cattle included in this study are presented in Table 1. Fourteen of the 17 cattle (83%) were treated for BRD within the first three weeks of the study, with a median time to first treatment of 5 days after arrival. Four of the 14 treated cattle (29%) were treated twice, with a median time to second treatment of 18 days after arrival; one of the 14 treated cattle (7%) was treated three times, with the third treatment given 27 days after arrival. By day 10 post-arrival, when CALA, TU, and TTW were completed for the first time, 12 of the 17 cattle had been treated for BRD, with 4 of those 12 cattle treated for the first time on day 10, when the sampling was done. By day 21 post-arrival, when CALA, TU, and TTW were completed the second time, 14 of the 17 cattle had been treated at least once for BRD, and two cattle had been treated twice. No mortality occurred.

**Table 1.** Descriptive data for the 17 cattle enrolled for transthoracic ultrasound (TU), computer-assisted lung auscultation (CALA), and transtracheal wash (TTW) cytology. Fourteen of the 17 cattle were treated for BRD once, 4 of the 14 treated cattle were treated a second time, and 1 of the 4 cattle treated twice was treated a third time (data not shown).

|  | Temp d 0<br>(° C) | First BRD<br>trt<br>(days)* | Second BRD<br>trt<br>(days)* | Weight<br>d 0<br>(kg) | Weight<br>d 12<br>(kg) | ADG<br>d 12<br>(kg) | Weight<br>d 26<br>(kg) | ADG<br>d 26<br>(kg) | Weight<br>d 82<br>(kg) | ADG<br>d 82<br>(kg) |
| --- | --- | --- | --- | --- | --- | --- | --- | --- | --- | --- |
| **Mean** | 39.4 | 5.9 | 20.8 | 225.4 | 230.5 | 0.4 | 240.6 | 0.6 | 285.5 | 0.7 |
| **St Dev** | 0.3 | 3.7 | 10.1 | 20.5 | 19.9 | 0.9 | 18.6 | 0.5 | 25.7 | 0.3 |
| **Median** | 39.4 | 5.0 | 18.0 | 230.9 | 238.6 | 0.5 | 239.1 | 0.6 | 293.6 | 0.7 |
| **Min** | 38.7 | 2.0 | 12.0 | 181.4 | 194.5 | -1.3 | 209.5 | -0.5 | 237.3 | 0.1 |
| **Max** | 39.9 | 16.0 | 35.0 | 263.6 | 259.5 | 1.6 | 269.5 | 1.5 | 344.1 | 1.3 |

\*Day of first and second BRD treatment is from arrival; for example, the mean day of first treatment was 5.9 days post-arrival.

### 3.2. Results of CALA, TU, and TTW

The initial study plan was to complete CALA, TU, and TTW for all 17 cattle on days 10 and 21 post-arrival. However, on day 10 it was evident that, due to logistical difficulties related to the portable handling facilities where the cattle were housed, it would not be possible to complete all tests on all 17 cattle in the time available. Therefore, on day 10, 16 cattle were assessed by CALA (one was too fractious to assess), and a subset of those 16 was assessed by TU (n = 10) and TTW (n = 9). Thus, on day 10 post-arrival, CALA, TU, and TTW results were fully obtained from nine of the 17 cattle. On day 21, we prioritized making all measurements from fewer cattle. Therefore, we completed CALA and TTW on 13 cattle and TU on ten of those 13 cattle. Four cattle were evaluated on both day 10 and day 21. Thus, combining the data collected on day 10 and day 21 yielded a complete set of CALA, TU, and TTW results for 18 measurements on 14 unique cattle, with four of those cattle being sampled on both day 10 and day 21.

The distribution of CALA scores measured on the right side of the thorax for both days is presented in Table 2. The CALA scores were relatively low, with 13 of 16 cattle assessed on day 10 post-arrival receiving a score of 1 or 2 and 11 of 12 cattle assessed on day 21 receiving a score of 1 or 2. No cattle received a CALA score of 4 or 5 on either day.

**Table 2.** Computer-assisted lung auscultation (CALA) and thoracic ultrasound (TU) scores from stocker cattle assessed 10 or 21 days after arrival (n = 12). Scores from CALA were assigned from 1-5 as defined by the computer’s algorithm, with 1=normal and 5=severely abnormal; scores from evaluation of the right side of the thorax are presented. No cattle received a CALA score of 4 or 5 on either day. Thoracic ultrasound (TU) scores for consolidation or comet tails were assigned by the examiner at the time of examination, with consolidation.

| Computer assisted lung auscultation score (n) | Day 10<br>(n = 16) | Day 21<br>(n = 12) |
| --- | --- | --- |
| Score = 1 | 3 | 5 |
| Score = 2 | 10 | 6 |
| Score = 3 | 3 | 1 |
| Thoracic ultrasound scores (n) | Day 10<br>(n = 10) | Day 21<br>(n = 10) |
| US consolidation $\geq 2$ cm<sup>2</sup> present on R or L side | 1 | 3 |
| US consolidation $\geq 5 \text{ cm}^2$ present on R or L side | 0 | 1 |
| Comet tail score R + L = 0 | 3 | 0 |
| Comet tail score R + L = 1 | 1 | 3 |
| Comet tail score R + L = 2 | 6 | 2 |
| Comet tail score R + L = 3 | 0 | 3 |
| Comet tail score R + L = 4 | 0 | 1 |
| Comet tail score R + L = 5 | 0 | 1 |
| Comet tail score R + L = 6 | 0 | 0 |

Ten cattle were evaluated by TU on days 10 and 21, and the results of TU scoring are presented in Table 2. Only one of eight cattle evaluated on day 10 had consolidation (>2 cm on one side). Only three of 10 cattle evaluated on day 21 had consolidation (two cattle had >2 cm of consolidation on one side, and one had >5 cm consolidation on one side). The calf with >5 cm of lung consolidation found on day 21 had been treated for BRD twice before day 21, once on day 5 post-arrival, and once on day 19 post-arrival. Comet tail artifacts were seen in most cattle but were generally not severe. On day 10, the highest score recorded (sum of scores on right and left sides of the thorax) was 2, found in 6 of 10 cattle examined. On day 21, the highest score recorded was a score of 5 (ID 152), recorded in 1 of 10 cattle examined, but most cattle (8 of 10) had a comet tail score of 2 or lower.

Nine and 13 cattle received TTW assessments on days 10 and 21, respectively; the results of TTW cytologic assessment are presented in Tables 3 and 4. Most cells found in the 200-cell differential counts were macrophages, neutrophils, and airway epithelial cells. On both days 10 and 21, the mean and median percent neutrophils were higher, and the mean and median percent airway epithelial cells and macrophages were lower in cattle previously treated for BRD compared to cattle not previously treated for BRD. Scores for mucus and blood were also higher in cattle previously treated for BRD, with maximum possible scores of 2 seen in TTW from cattle previously treated for BRD but not in cattle not previously treated for BRD. Four cattle received TTW on days 10 and 21: cattle 152, 178, 192, and 198. The percentage of TTW neutrophils was comparable at both time points for all 4 animals. On days 10 and 21, the percent TTW neutrophils were 64% and 65%, respectively, for calf 152; 63% and 66% for steer 192, and 63% and 85% for steer 198. The TTW from steer 178 was hypocellular, with no cells seen on day 10; on day 21, 1% neutrophils, 78% macrophages, 12% lymphocytes, 7% airway epithelial cells, and 2% multinucleated cells were seen. Steer 178 was one of 3 cattle never treated for BRD. All data for each animal sampled on days 10 and/or 21 are presented in Supplementary File S1.

**Table 3.** Results of the cytological assessment of transtracheal washes (TTW) collected from 9 stocker cattle 10 days after arrival, including results for all cattle, results for cattle not yet treated for BRD as of Day 10 (n = 3), and results for cattle treated for BRD before or on Day 10 (n = 6). AEC: airway epithelial cells; MP: macrophages; LYM: lymphocytes; NEU: neutrophils; NEU + Bac: neutrophils containing bacteria; EOS: eosinophils; MNC: multinucleated cells. No basophils or mast cells were seen in any sample at any time point. Mucus and blood were scored from 0 (little or no mucus or blood seen) to 2 (mucus or blood in most fields).

|  | AEC (%) | MP (%) | LYM (%) | NEU (%) | NEU + Bac (%) | EOS (%) | MNC (%) | Mucus (0 - 2) | Blood (0 - 2) |
| --- | --- | --- | --- | --- | --- | --- | --- | --- | --- |
| **Day 10, all cattle (n = 9)** |  |  |  |  |  |  |  |  |  |
| **Mean** | 8.2 | 51.3 | 0.7 | 26.8 | 0.1 | 0.1 | 2.1 | 0.7 | 1.0 |
| **Std Dev** | 7.5 | 27.2 | 0.7 | 26.9 | 0.3 | 0.2 | 2.4 | 0.8 | 0.9 |
| **Median** | 7.0 | 63.5 | 0.5 | 18.5 | 0.0 | 0.0 | 1.5 | 0.0 | 1.0 |
| **Min** | 0.0 | 0.0 | 0.0 | 0.0 | 0.0 | 0.0 | 0.0 | 0.0 | 0.0 |
| **Max** | 22.0 | 80.5 | 2.0 | 63.5 | 1.0 | 0.5 | 6.5 | 2.0 | 2.0 |
| Number of cattle with > 20% NEU: 4 of 9 |  |  |  |  |  |  |  |  |  |
| **Day 10, cattle not yet treated for BRD (n = 3)** |  |  |  |  |  |  |  |  |  |
| **Mean** | 13.9 | 49.0 | 0.8 | 0.3 | 0.0 | 0.0 | 2.7 | 0.3 | 0.7 |
| **Std Dev** | 9.9 | 34.6 | 0.8 | 0.5 | 0.0 | 0.0 | 2.5 | 0.5 | 0.9 |
| **Median** | 19.7 | 72.4 | 0.5 | 0.0 | 0.0 | 0.0 | 2.0 | 0.0 | 0.0 |
| **Min** | 0.0 | 0.0 | 0.0 | 0.0 | 0.0 | 0.0 | 0.0 | 0.0 | 0.0 |
| **Max** | 22.0 | 74.5 | 2.0 | 1.0 | 0.0 | 0.0 | 6.0 | 1.0 | 2.0 |
| Number of cattle with > 20% NEU: 0 of 3 |  |  |  |  |  |  |  |  |  |
| **Day 10, cattle previously treated for BRD (n = 6)** |  |  |  |  |  |  |  |  |  |
| **Mean** | 5.3 | 52.4 | 0.6 | 40.0 | 0.2 | 0.1 | 1.8 | 0.8 | 1.2 |
| **Std Dev** | 3.4 | 22.6 | 0.7 | 23.8 | 0.4 | 0.2 | 2.2 | 0.9 | 0.9 |
| **Median** | 5.0 | 49.0 | 0.3 | 44.6 | 0.0 | 0.0 | 1.3 | 0.5 | 1.5 |
| **Min** | 1.5 | 25.5 | 0.0 | 6.0 | 0.0 | 0.0 | 0.0 | 0.0 | 0.0 |
| **Max** | 11.0 | 80.5 | 1.5 | 63.5 | 1.0 | 0.5 | 6.5 | 2.0 | 2.0 |
| Number of cattle with > 20% NEU: 4 of 6 |  |  |  |  |  |  |  |  |  |

**Table 4.**
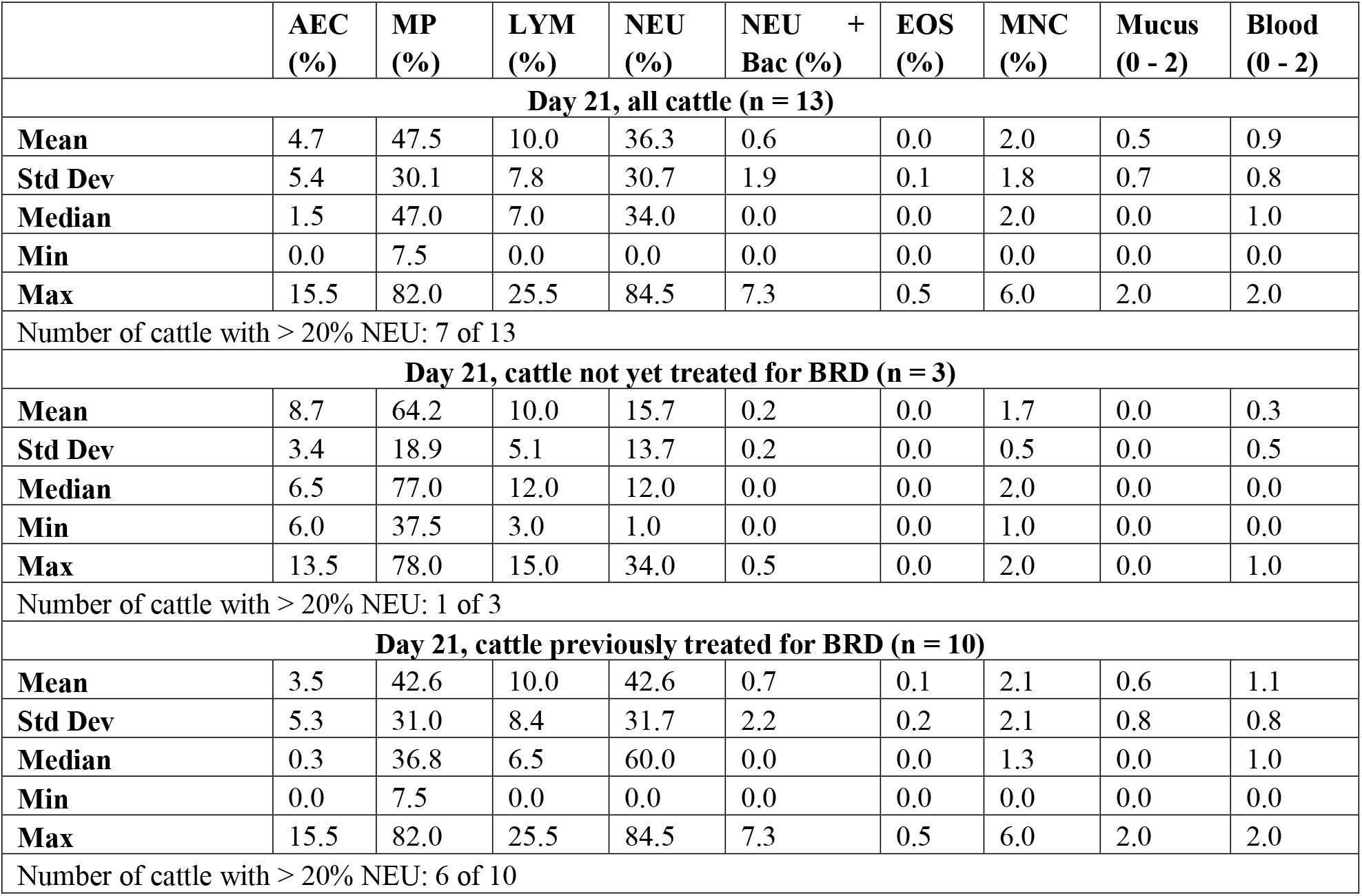
Results of the cytological assessment of transtracheal washes (TTW) collected from 12 stocker cattle 21 days after arrival, including results for all cattle, results for cattle not yet treated for BRD as of Day 21 (n = 3), and results for cattle treated for BRD before or on Day 21 (n = 9). AEC: airway epithelial cells; MP: macrophages; LYM: lymphocytes; NEU: neutrophils; NEU + Bac: neutrophils containing bacteria; EOS: eosinophils; MNC: multinucleated cells. No basophils or mast cells were seen in any sample at any time point. Mucus and blood were scored from 0 (little or no mucus or blood seen) to 2 (mucus or blood in most fields).

|  | AEC (%) | MP (%) | LYM (%) | NEU (%) | NEU + Bac (%) | EOS (%) | MNC (%) | Mucus (0 - 2) | Blood (0 - 2) |
| --- | --- | --- | --- | --- | --- | --- | --- | --- | --- |
| **Day 21, all cattle (n = 13)** |  |  |  |  |  |  |  |  |  |
| **Mean** | 4.7 | 47.5 | 10.0 | 36.3 | 0.6 | 0.0 | 2.0 | 0.5 | 0.9 |
| **Std Dev** | 5.4 | 30.1 | 7.8 | 30.7 | 1.9 | 0.1 | 1.8 | 0.7 | 0.8 |
| **Median** | 1.5 | 47.0 | 7.0 | 34.0 | 0.0 | 0.0 | 2.0 | 0.0 | 1.0 |
| **Min** | 0.0 | 7.5 | 0.0 | 0.0 | 0.0 | 0.0 | 0.0 | 0.0 | 0.0 |
| **Max** | 15.5 | 82.0 | 25.5 | 84.5 | 7.3 | 0.5 | 6.0 | 2.0 | 2.0 |
| Number of cattle with > 20% NEU: 7 of 13 |  |  |  |  |  |  |  |  |  |
| **Day 21, cattle not yet treated for BRD (n = 3)** |  |  |  |  |  |  |  |  |  |
| **Mean** | 8.7 | 64.2 | 10.0 | 15.7 | 0.2 | 0.0 | 1.7 | 0.0 | 0.3 |
| **Std Dev** | 3.4 | 18.9 | 5.1 | 13.7 | 0.2 | 0.0 | 0.5 | 0.0 | 0.5 |
| **Median** | 6.5 | 77.0 | 12.0 | 12.0 | 0.0 | 0.0 | 2.0 | 0.0 | 0.0 |
| **Min** | 6.0 | 37.5 | 3.0 | 1.0 | 0.0 | 0.0 | 1.0 | 0.0 | 0.0 |
| **Max** | 13.5 | 78.0 | 15.0 | 34.0 | 0.5 | 0.0 | 2.0 | 0.0 | 1.0 |
| Number of cattle with > 20% NEU: 1 of 3 |  |  |  |  |  |  |  |  |  |
| **Day 21, cattle previously treated for BRD (n = 10)** |  |  |  |  |  |  |  |  |  |
| **Mean** | 3.5 | 42.6 | 10.0 | 42.6 | 0.7 | 0.1 | 2.1 | 0.6 | 1.1 |
| **Std Dev** | 5.3 | 31.0 | 8.4 | 31.7 | 2.2 | 0.2 | 2.1 | 0.8 | 0.8 |
| **Median** | 0.3 | 36.8 | 6.5 | 60.0 | 0.0 | 0.0 | 1.3 | 0.0 | 1.0 |
| **Min** | 0.0 | 7.5 | 0.0 | 0.0 | 0.0 | 0.0 | 0.0 | 0.0 | 0.0 |
| **Max** | 15.5 | 82.0 | 25.5 | 84.5 | 7.3 | 0.5 | 6.0 | 2.0 | 2.0 |
| Number of cattle with > 20% NEU: 6 of 10 |  |  |  |  |  |  |  |  |  |

### 3.3. Data analysis

Too few cattle (day 10, n=1; day 21, n=3) had lung consolidation identified by TU to analyze. There was no significant relationship between TU comet tail score and TTW inflammation (P = 0.8385); between TTW inflammation and CALA score on the right side of the thorax (P = 0.1445), the left side (P = 0.8138), or the sum of the measurements on the right and left sides (P = 0.6551); or between TU comet tail score on the right or left, and the CALA score on the right, left, or sum of right and left (P-values ranged from 0.3078 to 1.0000). The relationship between a clinical diagnosis of BRD at any time point and CALA score on the right (P = 0.8519), the left (P = 0.8021), or the sum of the right and left (P = 0.9922) was not significant; nor was the relationship between clinical BRD diagnosis and comet tail score (P = 0.7264). The relationship between a clinical diagnosis of BRD and TTW inflammation on day 10 or 21 was statistically significant (χ^2^ = 3.5762; P = 0.0586); because of this, the relationship between clinical diagnosis of BRD on 0, 1, 2, 3, 4, 5, 6, or 7 days within the identification of TTW inflammation on day 10 or day 21 was also evaluated. The relationship between clinical BRD diagnosis and TTW inflammation was statistically significant when a clinical BRD diagnosis was made within two days of finding an inflammatory TTW (χ^2^ = 7.9050; P = 0.0702); 4 of 4 cattle had an inflammatory TTW when diagnosed with BRD within two days, while only 6 of 17 cattle had an inflammatory TTW when not diagnosed with BRD within 2 days.

## 4. Discussion

The lack of an accurate method of BRD diagnosis that can be rapidly applied to large numbers of animals in the field remains a problem that impairs the ability of producers, veterinarians, and researchers to treat, control, and prevent BRD. In recent years, TU has been evaluated by multiple research groups looking for a reliable and practical method to diagnose BRD in the field. In dairy calves, TU has been used to identify lung lesions associated with important outcomes such as weight gain, time to first pregnancy, and milk produced in the first lactation [5-7]. In feedlot cattle, the maximal depth and area of lung consolidation identified by TU were significantly associated with an increased risk of mortality or relapse after the first treatment [8,12]. However, TU to identify lung consolidation still lacks sensitivity for BRD diagnosis in feedlot cattle. When various cutoffs of depth and area of consolidation were evaluated in cattle diagnosed with BRD by a combination of clinical signs and culture of high numbers of *Mannheimia haemolytica* from airway lavage, the best combination of sensitivity and specificity was found at consolidation depth of > 5 cm, which yielded 75% sensitivity (95% CI: 43-95%) and 82% specificity (95% CI: 57-96%) [12]. The sensitivity of TU in feedlot cattle may be diminished by the fact that ultrasound of the right cranial lung lobe, where pneumonic lesions often occur, is not feasible in feedlot cattle due to their size and the constraints of access in chutes used for restraint.

Stethoscope auscultation is a vital diagnostic procedure in evaluating active airway disease within veterinary clinical settings [21,24]. However, subtle abnormalities in airway sounds can be challenging to detect and remain semi-objective in severity determination. As such, CALA systems provide relative standardization to overcome such limitations. Recent studies have utilized this technology with encouraging results, albeit in relatively small trials; two studies determined that computer-assisted auscultation systems resulted in BRD diagnostic sensitivities and specificities of 88-93% and 75-90%, respectively [9,10]. Recently, CALA was used to identify feedlot cattle most likely to benefit from metaphylactic antimicrobial treatment [11].

Cytological evaluation of the TTW to identify lower airway inflammation remains understudied in bovine medicine. TTW, the collection of tracheal secretions via tracheal perforation followed by instillation then aspiration of an aliquot of sterile physiologic saline, has been used for decades to identify cytologic evidence of lower airway inflammation in horses [15,25]. Cytologic assessment of the proportion of cell types, including airway epithelial cells, neutrophils, macrophages, lymphocytes, and eosinophils, is used to determine the type of airway inflammation present in the sampled patient. Cytologic assessment of TTW from horses with no evidence of lung disease has been reported to reveal a majority of airway epithelial cells, macrophages, and lymphocytes, with fewer neutrophils [16]. While the range of percent TTW neutrophils found in apparently normal horses has been reported to overlap with the range found in horses with various types of lower airway disease [26,27], an increased proportion of neutrophils relative to other cell types has repeatedly been identified in TTW from horses with pneumonia or suppurative bronchitis [25,26].

In cattle, TTW has mainly been used to identify infectious agents in the lower airways while bypassing contamination from upper respiratory commensals [14,28,29]. In contrast, bronchoalveolar lavage (BAL) has more often been used to identify cytologic evidence of lower airway inflammation [30,31]. Allen et al. compared cytologic findings of BAL fluid collected from the right cranioventral lung by endoscopic BAL from [59] feedlot cattle with clinical BRD and 60 clinically normal controls in the same feedlot [30]. This investigation revealed that neutrophils were higher and macrophages were lower in BAL from BRD cases relative to controls, with statistically significant differences in cell populations between the groups.

However, some case animals had cytologic differential counts that appeared normal, and some controls had differential counts that suggested an inflammatory response. The authors acknowledged that these discrepancies may have been due to failure to lavage abnormal lung in some BRD cases, inaccurate clinical diagnosis, or airway inflammation that did not necessarily represent disease.

A TTW provides a composite sample of airway secretions moving up from all bronchi, bronchioles, and alveoli. At the same time, the BAL represents a sample of the single bronchoalveolar unit into which the tube used for BAL collection is wedged. Therefore, a TTW should provide a more representative sample than a BAL for lung disease that is focal or regional and not generalized, as can be the case in BRD. Although the TTW provides a composite sample of the lower airways, it may also be influenced by tracheal or even laryngeal or pharyngeal inflammation. Cytologic characterization of the TTW in cattle with inflammation limited to the pharynx, larynx, or trachea has not, to our knowledge, been reported.

In the present study, because normal cytologic findings of the bovine TTW have not been defined, we chose a cutoff of 20% neutrophils or greater to define an inflammatory response based on criteria described for identifying airway inflammation in horses with heaves using BAL [23]. While BRD and heaves are diseases with entirely different etiologies, we assumed that the proportion of neutrophils found in an airway wash from either a healthy cow or horse would be similar (i.e., containing approximately less than 20% neutrophils). The fact that our analysis revealed that airway inflammation, as defined in this study, was the test result most closely related to the clinical diagnosis of BRD suggests that cytologic evidence of inflammation in the TTW warrants further investigation in research aiming to accurately identify the antemortem presence of respiratory disease in cattle. This is made apparent by our finding that all cattle diagnosed with BRD within two days possessed an inflammatory TTW.

The lack of a statistically significant relationship between the results of the three tests evaluated, CALA, TU, and TTW was somewhat surprising. We expected a closer relationship between these tests, and our findings suggest that these tests measure different and possibly unrelated outcomes. However, one limitation of this study was that most of the cattle were not sampled on the day their clinical diagnosis of BRD was made; this may have contributed to the lack of a significant relationship between the tests evaluated. The small number of cattle sampled is another significant limitation of this study, which may have increased type II statistical error.

To our knowledge, this study is the first of its kind and was aimed at determining which diagnostic tests might be feasible for use in future more extensive trials of methods to prevent or control BRD in stocker cattle and the relationship between the results of those tests. Although the time required to collect a TTW rules out the use of such a test for real-time diagnosis of BRD in commercial stocker operations, the relationship identified between TTW inflammation and recent treatment for BRD suggests that cytologic assessment of airway inflammation via TTW merits further investigation as a method to help confirm antemortem respiratory disease in research to develop practical and accurate methods to diagnose BRD.

## 5. Conclusions

This pilot study demonstrated little association with CALA, TU, and TTW testing outcomes, which may indicate their independent associations with disease outcomes. Through independent analyses, we identified that the assessment of TTW cytology, utilizing >20% neutrophil concentration to demark airway inflammation, was significantly associated with clinical diagnosis of BRD, and was significantly associated with a BRD diagnosis within two days of sampling. While the logistical feasibility of performing TTW in a commercial setting is questionable, these results provide a promising method of improving antemortem diagnosis of BRD in controlled settings and warrants further investigation in larger, more diverse studies. Supplementary Materials: Table S1: Treatment, CALA, TU, and TTW cytology scores for cattle on days 10 and 21.

## Author Contributions

Conceptualization, MAS and ARW; methodology, MAS, ARW, HFC, and CB; software, RWW; validation, MAS, HFW, and RWW; formal analysis, MAS and RWW; investigation, MAS, ARW, HFC, and BBK; resources, ARW and BBK; data curation, ARW and HFC; writing—original draft preparation, MAS and ARW.; writing—review and editing, MAS, ARW, HFC, RWW, CB, and BBK; visualization, MAS, ARW, and RWW; supervision, ARW and BBK; project administration, MAS, ARW, and BBK. All authors have read and agreed to the published version of the manuscript.

## Institutional Review Board Statement

The animal study protocol was approved by the Animal Care and Use Committee of Mississippi State University (protocol #17-120) and carried out in accordance with relevant IACUC and agency guidelines and regulations.

## Acknowledgments

The authors gratefully acknowledge the assistance provided by R. Tucker Wagner and the staff of the Mississippi State University College of Veterinary Medicine (MSU CVM) Clinical Pathology Laboratory. The authors also thank students and staff of the Mississippi Agri-cultural and Forestry Experiment Station (MAFES) and Mississippi State University College of Veterinary Medicine for their assistance in animal care and sample collection. The Department of Pathobiology and Population Medicine at the MSU-CVM and Department of Large Animal Clinical Sciences at the TAMU SVMBS provided internal financial support for the study.

